# Spatio-temporal variability of airborne viruses over sub-Antarctic South Georgia: Influence of sampling methodology and proximity of the ocean

**DOI:** 10.1101/2024.08.23.609320

**Authors:** Ritam Das, Lucie Malard, David A. Pearce, Peter Convey, Janina Rahlff

## Abstract

Studying airborne viruses in remote environments like the sub-Antarctic island South Georgia offers key insights into viral ecology, diversity, and their role in shaping ecosystems through microbial and nutrient interactions. We analyzed airborne viral community composition at two sites in South Georgia. Sampling took place using multiple methodologies, with the data produced subjected to viral metagenomics. The Coriolis µ device (wet collection) was the most effective, yielding 30 viral scaffolds. Two-thirds of the scaffolds were only obtained from the coastal location, indicating that location influences airborne viral diversity. Protein-based clustering of 39 viral operational taxonomic units (vOTUs) revealed similarities of 15 with known marine viruses, suggesting oceanic influence on the airborne viral community. Protein homologs related to UV damage protection and photosynthesis from two airborne vOTUs were widely distributed across major oceans, suggesting their potential role in supporting the resilience of marine microorganisms under changing climate conditions. Some vOTUs had protein similarities to viruses infecting extremophiles, indicating viral adaptations to harsh environments. This study provides a baseline for understanding the complexity and sustainability of airborne viral communities in remote ecosystems. It underscores the need for continued monitoring to assess how these communities respond to shifting atmospheric and ecological conditions.

## Introduction

The cryosphere, encompassing ice sheets, glaciers, permafrost, and sea ice, covers approximately 20% of Earth’s surface, and hosts some of the planet’s most climatically sensitive ecosystems [1]. Characterized by chronically low temperatures and harsh environmental conditions and covered by continental-scale ice sheets on average 2 km thick, the fifth-largest continent Antarctica is a critical driver of global climate and ocean systems. Its final separation from southern South America and Australia around 33 million years ago in the final stages of the breakup of Gondwana was followed by cooling and the eventual formation of its ice sheets, and led to the evolution of a diverse cold-adapted microbiota [2]. Aerial dispersal is a major route for the invasion of new species [3], particularly those that are small enough, or have suitable propagules, to be uplifted into and transported into the air column. Antarctica is increasingly appreciated to host a complex microbial diversity [4, 5], much of which appears to be endemic to the continent [6, 7]. However, few studies have yet investigated airborne bacteria in the Antarctic region [8–11]. Furthermore, there is a notable scarcity of research on both Antarctic viral diversity generally and on airborne viruses specifically (reviewed by Heinrichs, et al. [12]), which is rather astonishing considering that these biological entities are ubiquitous and reach global numbers of 10^31^ viral particles [13]. One reason for the absence of viral investigations is that DNA yields from aerosols are usually low due to the very limited biomass, which precludes metagenomic studies that could otherwise identify viruses. Instead, 16S rRNA gene surveys (metabarcoding) involving a PCR step for amplification are often applied [14, 15], which are appropriate for studies of prokaryotes. Viruses typically do not disperse in the air as free particles; instead, they often attach to soil dust or marine organic aggregates, which can then form aerosols [16]. The deposition rates of viruses in the atmosphere, which can be up to ∼460 times higher than those of bacteria, are positively correlated with organic aerosols [16]. This suggests that viruses may have longer residence times in the atmosphere than microbes or other microscopic propagules, likely due to their smaller size [17]. Consequently, there is greater likelihood of their dispersal over long distances.

Sea spray aerosols (SSA), which are formed by the primary emission from the ocean, are the key component by mass of marine aerosols and play a crucial role in the Earth’s climate system [18]. A virus infecting the marine coccolithophore *Emiliania huxleyi* can lead to their rapid demise and induce coccolith shedding, the process by which these marine algae release their calcium carbonate plates into the surrounding water. This can lead to subsequent incorporation of the calcite units (coccoliths) to the SSA, with these viruses acting as a regulator of the mass of marine aerosols [19]. Through aerosol-mediated long-distance transport, viruses could constitute an ecological and evolutionary driving force beyond marine ecosystems alone, for instance through rain-washout events, where aerosolized particles are removed from the atmosphere by precipitation, or through direct sedimentation [16]. Such events would allow airborne viruses to interact with microbes and other organisms in terrestrial ecosystems, affecting their abundance, community composition and distribution in the recipient ecosystems [16].

Investigations of viral diversity and lifestyle have been conducted on microbial viruses from Antarctica’s lakes [20], sea ice [21, 22], under the ice shelf [23], soils [24], from cryoconite holes [25] and marine ecosystems including the Southern Ocean [26, 27]. In a study of Antarctic surface snow, similar viral populations were described from locations separated by several hundred to thousands of kilometres and overlaps between snow and marine viral populations were apparent [28]. Flavobacteria from Antarctic snow carry adaptive immunity in the form of clustered regularly interspaced short palindromic repeats (CRISPR)-Cas systems, indicating that defence mechanisms against mobile genetic elements are present [29]. Recent work has demonstrated that viral diversity in the Southern Ocean around the Western Antarctic Peninsula is complex and has uncovered both diversity and novelty amongst the viruses present. As well as tailed dsDNA bacteriophages representing the class Caudoviricetes, abundant eukaryotic viruses including polinton-like viruses, nucleocytoplasmic large DNA viruses (NCLDV), and virophages have been reported [30]. A study in the Antarctic and sub-Antarctic region of South Georgia observed the mortality and morbidity of several key indigenous species due to the spread of a high pathogenicity avian influenza virus H5N1 [31]. Therefore, monitoring viruses in isolated (including anthropogenic) locations could help track changes in viral communities over time, determine their stability and resilience, and assess the importance of anthropogenic influence. Furthermore, contemporary increase in global temperature and its regional impact on cryosphere-associated life forms presents a unique opportunity to understand ecosystem function, which could help shape a sustainable ‘One World’ for the future.

In this study, we used viral metagenomics to investigate the spatio-temporal variability of airborne viruses on the remote sub-Antarctic island of South Georgia in the South Atlantic Ocean, collected over seven days using different air samplers and a rain collector. We hypothesised that the surrounding ocean would be a major contributing source for atmospheric viruses in the bioaerosols but also sought homologs of viral proteins obtained in our South Georgia samples in datasets available from other oceans to which viruses are probably dispersed or from which they are recruited.

## Materials and methods

### Field sampling and filtration

Air and rain samples were collected in South Georgia (latitude −54.4, longitude −36.5), which is ∼1800 km from Antarctica to the south, ∼1400 km from the tip of the Antarctic Peninsula to the south-west and ∼2150 km from southern South America to the west. Collection took place from October 15 to 21, 2021, at two different sampling sites: King Edward Point (KEP) and Deadman’s Cairn at Lewis Pass (DMP) (Figure 1). These sites represent two different sampling altitudes of ∼3 and ∼200 m above sea level, respectively. Site KEP (−54.283528, −36.495194) was directly at the coast, while DMP was located 2 km inland (−54.269964, −36.503672, Figure 1f). Three different air samplers were used, namely the “wet” Coriolis ® µ air sampler, “dry” Coriolis Compact air sampler (both Bertin Technologies SAS, Montigny-le-Bretonneux, France), and a Whatman Vacuum Pump attached to a Sartorius 250 ml filter holder collector (“pump”). The samplers were positioned about 1 m above the ground surface. For rain sampling, a plastic funnel attached to a 1 L sterile collection bottle (“collector”) was used (at KEP only). At the DMP site, only the wet and dry Coriolis samplers could be used as they function on batteries and do not require main power. Sampling duration was 1-6 h when using the Coriolis instruments with high flow rates, whether in wet or dry conditions, and extended up to 22 h when air was collected using the vacuum pump. Rain collectors were used for 15 h (Table S1). We extracted hourly wind speed, wind direction, pressure, and air temperature data for the sampling period (Table S1) from the Grytviken Automatic Weather Station. For each sample, the corresponding values of each variable were averaged to cover each sampling duration, allowing us to obtain average environmental conditions for each sample.

**Figure 1:**
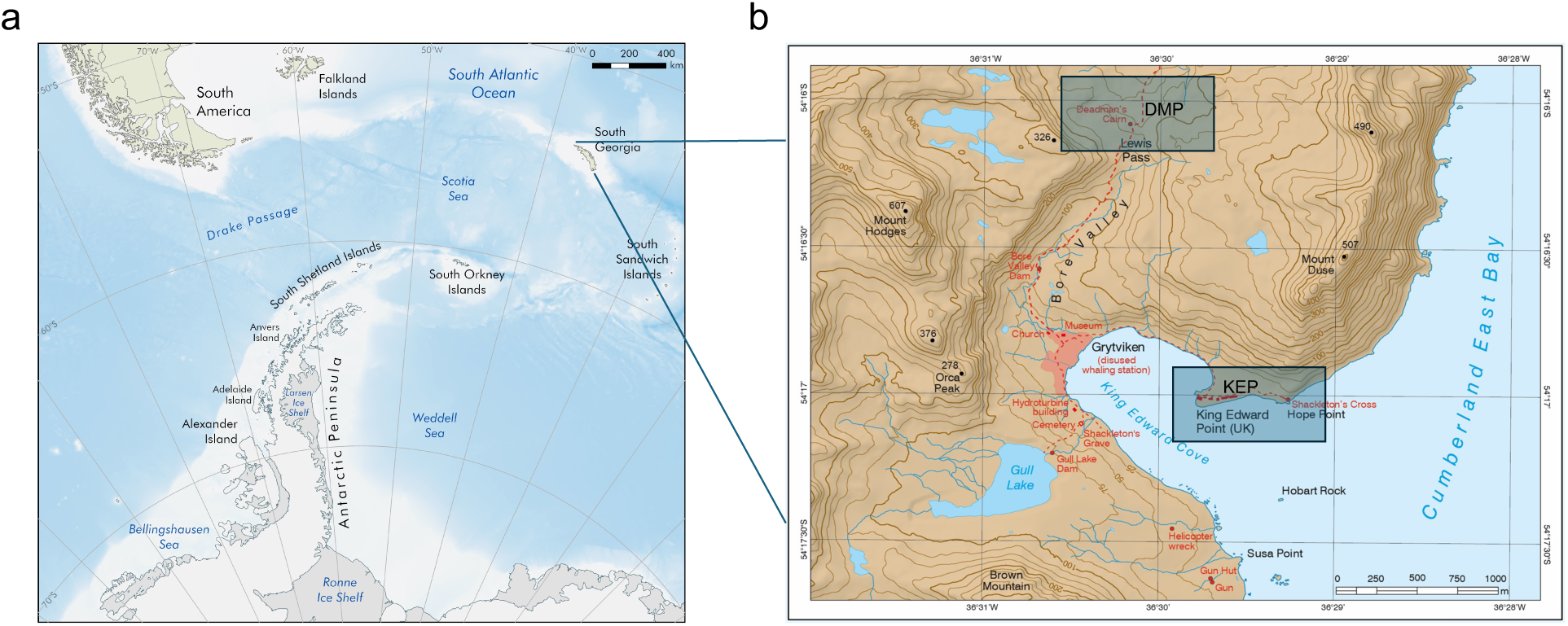
Map showing the location of South Georgia (a) and the two sampling sites DMP = Deadman’s Cairn, Lewis Pass, and KEP = King Edward Point (b). Sources: a) Mapping and Geographic Information Centre, British Antarctic Survey, 2025; Bathymetry: Weatherall, et al. [114], Coastline: SCAR Antarctic Digital Database, 2024 for areas south of 60°S and Natural Earth for areas north of 60°S b) South Georgia and The Shackleton Crossing” by the British Antarctic Survey, 2021, 1:200,000 and 1:40,000 scale, BAS (Misc 12), Cambridge, British Antarctic Survey. Reproduced with permission.

### DNA extraction and sequencing

Cellulose nitrate membrane filters with air samples were sliced into 0.5 mm wide strips using a sterilized scalpel to maximize the filters’ exposure to the cell lysis buffer during DNA extraction. For samples collected in nuclease-free water using the Coriolis wet device, the collection cones were vortexed (SciQuip, Shropshire, UK) before being filtered through 0.2 μm pore size cellulose nitrate membrane filters (GE Healthcare Life Sciences, Chicago, IL, USA) using a Welch WOBL vacuum pump (Welch, Mt. Prospect, IL, USA) for 10 min. Samples obtained via the Coriolis Compact sampler (the dry device) were resuspended in 15 mL of DNA/RNAase-free water, manually shaken, and vortexed before undergoing the same filtration process. All pre-processing was conducted in a Class II cabinet, and the vacuum pump, tubing, and filtration apparatus were sterilized with 70% ethanol before use. DNA extraction from air samples was performed using the Qiagen PowerSoil kit (Qiagen, Hilden, Germany) according to the manufacturer’s instructions. Sterile water controls taken during sampling and extraction kit controls, although not sequenced for metagenomics, showed no contamination in the 16S rRNA gene amplicon dataset, as detailed in a separate, currently submitted manuscript [32]. DNA extracts were submitted to NU-OMICS sequencing facility at Northumbria University, Newcastle, for Illumina NextSeq shotgun metagenomics sequencing. DNA libraries were prepared using the Nextera XT DNA Library Prep Kit, following the manufacturer’s instructions. Sequencing was conducted on an Illumina NextSeq system (Illumina Inc., USA) with V2.5 300-cycle chemistry.

### Metagenomic analysis

Sequencing reads went through adapter trimming using BBduk within BBTools [33], and afterwards Sickle v.1.33 [34] was run in paired-end mode and ‘-t sanger’ setting. Taxonomic profiling of microbes was performed using the tool mOTUs v.3.0.2 on trimmed reads with options ‘-A’ (reports full taxonomy) ‘-c’ (reports counts) ‘-M’ (to save intermediate marker gene cluster count) including a separate run to retrieve unassigned taxa. The tool mOTUs employs universal, protein-coding, and single-copy phylogenetic marker gene sequences [35, 36]. The trimmed reads were subsequently assembled using metaSPAdes v.3.15.5 [37] and metaviral SPAdes [38]. Assemblies were combined and viral scaffolds were predicted by running VIBRANT v.1.2.1 [39], VirSorter v2 with setting ‘--include-groups “dsDNAphage,ssDNA,NCLDV,lavidaviridae“’ and default score [40], and geNomad v.1.3 [41]. Viral scaffolds detected by the different tools were combined and filtered to a minimum length of 3000 bp. The number of assembled viral scaffolds by sampling device was summed. As the number of viruses and the genome size were very low, assemblies across all read files (respective forward and reverse reads of all samples merged) were performed using the above tools and additionally MEGAHIT v.1.2.9 [42]. The resulting viral scaffolds were combined with the previous ones, and a size cut-off of 5000 bp was applied. Viruses were clustered using VIRIDIC v.1.0 r3.6 [43], and only one viral operational taxonomic unit (vOTU) of each species cluster (demarcation threshold = 95 % intergenomic similarity) was used in downstream analysis, and CheckV v.1.0.1 [44] was run. To determine if a vOTU was present in a sample, reads were mapped back to vOTUs using Bowtie 2 v.2.3.5.1 [45] with settings ‘--mp 1,1 --np 1 --rdg 0,1 --rfg 0,1 --score-min L,0,-0.1’ [46], which ensure that only reads with 90% sequence identity map to the vOTU. To follow guidelines of Roux, et al. [47], a vOTU was considered present if at least 75% of the scaffold was covered with reads, which was checked using the ‘calcopo.rb’ script [28]. Mean coverages of vOTU were determined using the ‘04_01calc_coverage_v3.rb’ script [48] and coverages were normalized to sequencing depth. Relative abundance was calculated as the proportion of each normalized coverage relative to the total, expressed as a percentage. Gene calling was performed using Prodigal v.2.6.3 [49], and for viral clustering, vConTACT2 v.0.9.19 [50] was run on the vOTU proteins together with those from the 1August2023 viral reference database (https://github.com/RyanCook94/inphared/tree/a330daa635cd3c78843d470668cb22ff842960e4) derived from INPHARED [51]. Results were compiled using graphanalyzer v.1.5.1 [52] and a network was built in Cytoscape v.3.10.3 [53]. PhaMer [54] and PhaGCN [55] within PhaBOX v.1 [56] were used to discriminate phages from non-phages, and to determine the taxonomic rank of a phage at family level, respectively. VirClust webtool [57] with default settings was run on the airborne vOTUs. ViPTree v.4.0 [58] was run with settings for dsDNA for nucleic acid type of reference viruses and “Any host” as the host category. Further virophage classification was performed using https://github.com/simroux/ICTV_VirophageSG [59]. In a previous, non-polar study, viral genomes retrieved from aerosols and rainwater had a significantly higher guanine/cytosine (GC) base content compared to marine viruses [60], which is why the GC content of the Antarctic air vOTUs was further explored. The GC content of vOTUs was determined using EMBOSS v. 5.0.0 [61]. Annotations of vOTUs were performed using DRAM-v v.1.4.6. [62], which includes annotations from the databases Pfam, VOGDB, KOfam, dbCAN, and RefSeq viral. The vOTUs were BLASTed against the IMG/VR Viral Nucleotide Database v.4 with an e-value cut-off of 1e-5 [63]. Domain analysis of the proteins was performed using NCBI Conserved Domain Search website [64]. A BLASTp analysis for selected amino acids sequences from two vOTUs was run against the Ocean Gene Atlas [65, 66] using the Tara Oceans Microbiome Reference Genome Catalog v1 OM-RGC_v1 (based on metagenomes) and an e-value cut-off of 1e-10. Photosynthetically active radiation (PAR) was chosen as the environmental variable based on which the spatial distribution and abundance (average copies per cell) of the protein homologs of vOTU_35_ORF14 and vOTU_39_ORF2 was explored, because the protein homologs are functionally related to PAR or ultraviolet radiation, which both correlate with global solar radiation [67]. The phylogenetic tree acquired from the Ocean Gene Atlas was edited using the Interactive Tree of Life (iTOL) v.6 website [68]. Overlaps of vOTUs between stations were visualized with Venn Diagrams made with the Ugent webtool (https://bioinformatics.psb.ugent.be/webtools/Venn/). Host predictions were performed using iPHoP v.1.2.0 and the database Sept_2021_pub [69].

### Diversity analysis and statistics

Community and diversity analysis of vOTUs was performed using phyloseq v. 1.48.0 [70] within the R programming environment v.4.4.0 [71]. Plotting and statistical analysis of Shannon-Wiener index was performed in Graphpad Prism 10.

## Results

### Viral community composition is affected by sampling site, collection device, and daily fluctuations

The different sampling devices collected different numbers of viral genome fragments. From the 60 assembled vOTUs >3 kb length, most vOTUs (30) were assembled from samples collected with the wet Coriolis and only one vOTU from the dry Coriolis (Figure 2a, collection devices depicted in Figure 2c). Seven and 22 vOTUs were assembled after collection with the vacuum pump and the rain collector, respectively. Of the 60 assembled vOTUs, 43 and 17 vOTUs were collected at the coastal KEP site and the inland higher altitude DMP site, respectively (Figure 2d). From these assembled scaffolds and the cross-assembled ones, 39 vOTUs >5 kb were identified and further analyzed. Mapping reads to these vOTUs showed that 15 unique vOTUs were present in samples taken with the rain collector, another unique 15 vOTUs in samples from the wet Coriolis, and 9 vOTUs occurred in samples obtained by both devices, suggesting that these two sampling methods complement each other for virus sampling (Figure 2b). Eleven vOTUs were shared between both sampling sites, while 19 were only present at KEP and 9 at DMP (Figure 2e).

**Figure 2:**
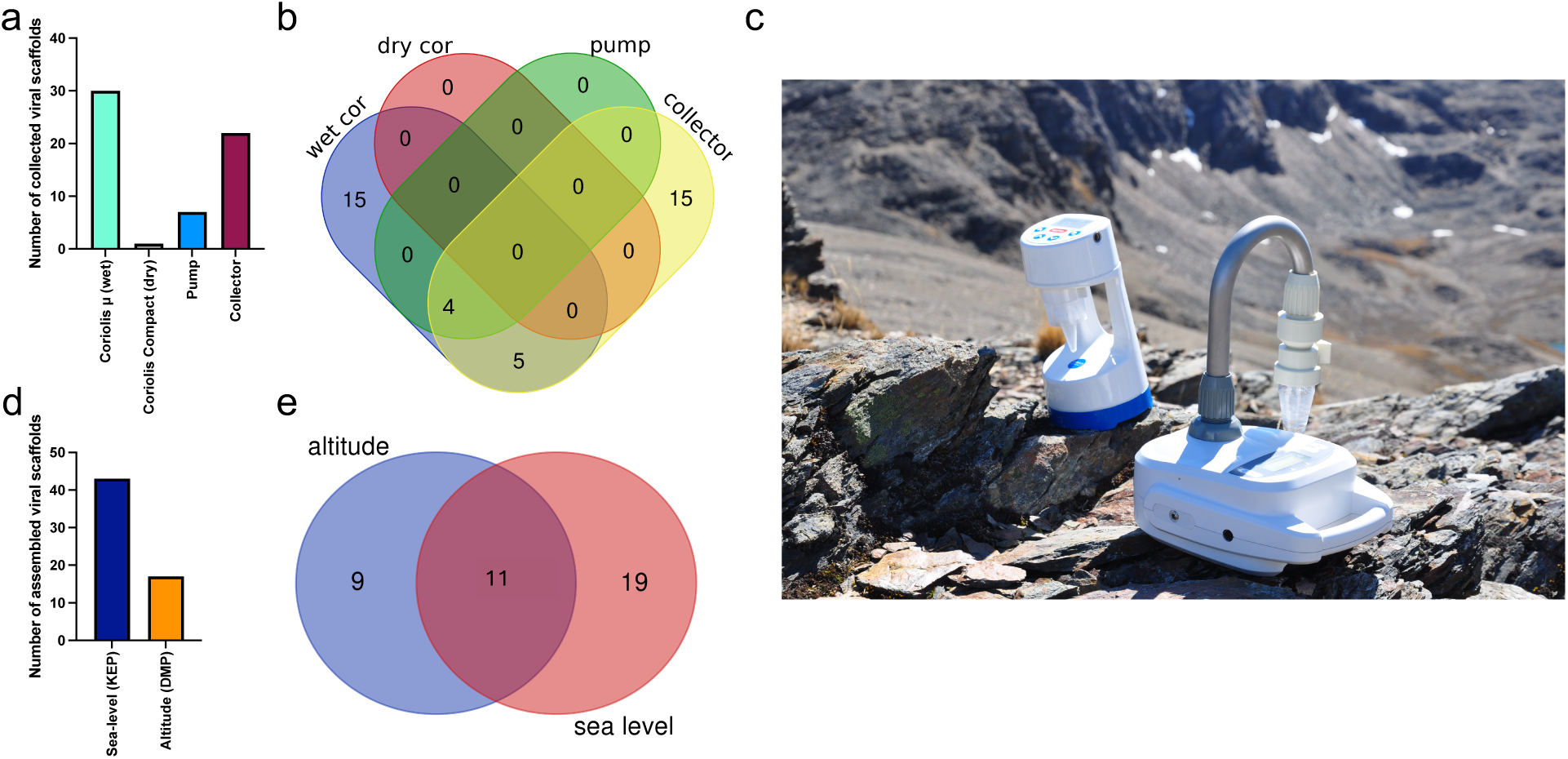
Airborne virus sampling with different collection devices from two sampling sites in South Georgia. a) Number of collected viral scaffolds >3000 bp per sampling device; for description of sampling devices, see main text. b) Venn diagram showing overlapping vOTUs for different sampling devices based on read mapping. c) Coriolis µ and Coriolis COMPACT sampler in the Antarctic landscape (Image: David Pearce). d) Number of collected viral scaffolds >2 kb by sampling site, e) Venn diagram showing overlapping vOTUs (>5 kb) between sampling sites based on read mapping.

The vOTUs were clustered into viral clusters (VCs) by vConTACT2 corresponding to genus level clusters, while “singletons” and “outliers” may represent novel viruses. Relative abundance analysis showed that VC_1229_0 (containing vOTU_12 & vOTU_13), vOTU_24, and vOTU_30 were the most abundant (based on coverage depth) and prevalent (based on coverage breadth) VCs in the community (Figure 3a, Table S2). Based on Virsorter2, VC_1229_0 might contain virophage vOTUs, family Lavidaviridae, which could however not be further validated by using virophage-specific markers. VOTU_30 was classified as Bamfordvirae by geNomad, which includes NCLDV. Shannon-Wiener diversity was highest for vOTUs sampled with the rain collector at KEP (2.9 and 3.0) compared to the wet Coriolis at DMP (2.1 and 2.3) and compared to the wet Coriolis and pump at KEP (0.6 – 1.9) and thus did not show a clear Shannon-Wiener diversity difference between the two sites but may indicate a joint effect of the sampling device and site on alpha-diversity (Figure 3b).

**Figure 3:**
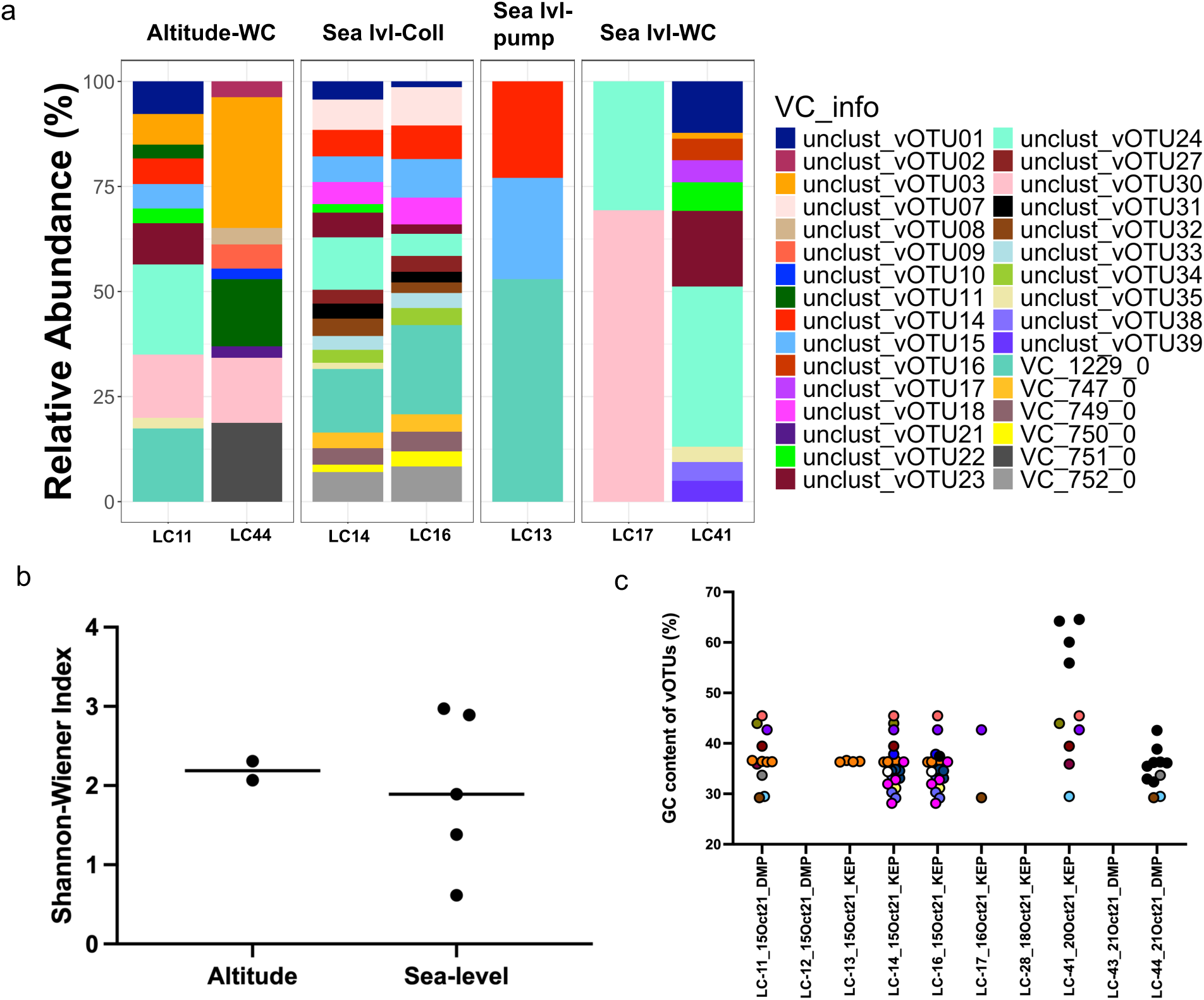
Viral cluster and vOTU distribution and relation to guanine/cytosine (GC) content. a) Relative abundance of viral clusters (VC) and unclustered viral operational taxonomic units (vOTUs) across stations. b) Shannon-Wiener index for viruses separated by study site. c) GC content of vOTUs versus sampling date and site. If vOTUs were found in several samples based on read mapping, they have the same color; if they are unique for a sample, they are black. DMP = Deadman’s Pass, KEP = King Edward Point, Sea/lvl = Sea-level, WC = wet Coriolis, LC refers to internal sample identification.

During sampling on October 15, 2021, both the vacuum pump and the collector captured some of the most abundant vOTUs (Figure 3a). Based on read mapping, vOTUs and VCs reoccurred and co-occurred on different days and at both sites, but also new vOTUs appeared (Figure 3a & c). Rain samples (LC-14 & LC-16) contained some vOTUs that were also recovered from air on October 15, suggesting that these overlapping vOTUs were washed out by falling rain from the atmosphere or resulted from raindrops that were collected with the aerosol sampler. Some VCs were only detected in the rainwater, namely VC_752_0, VC_750_0, VC_749_0 and VC_747_0 (all predicted Caudoviricetes). Notably, on one day (LC-41), where viruses were sampled at KEP with the wet Coriolis, four vOTUs present in the sample based on read mapping had a GC content of 61.8% ± 4.0 (mean ± standard deviation, Figure 3c, Table S3).

### Protein-based clustering reveals similarities to known marine viruses

According to classification of the 39 vOTUs by geNomad, Adintoviridae (3), Caudoviricetes (15 including Crassvirales (3)), Imitervirales (1), Bamfordvirae (2), unclassified (10) or not determined by the tool (8) were found. VirSorter2 predicted dsDNA phages (13), Lavidaviridae (9), NCLDV (7) and not determined (10) (Table S3). Depending on the virus identification tool, evidence for the presence of 13 – 17 phages among the 39 vOTUs was found (Figure 4a, b, Table S3). Four phages were predicted to be family Demerecviridae and two others family Straboviridae (Table S3).

**Figure 4:**
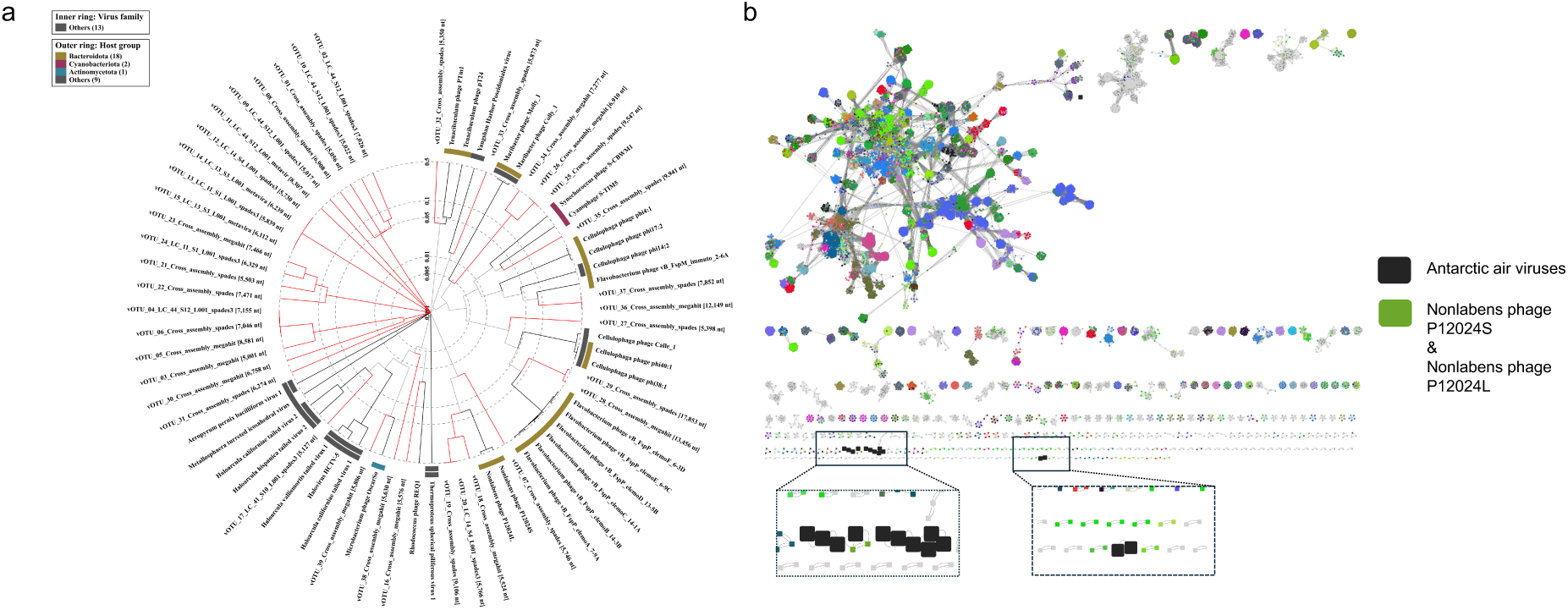
Viral protein-based clustering. a) Proteomic tree of 39 viral operational taxonomic units (red branches) and their related viruses constructed using ViPTree. b) Protein-sharing network of Antarctic vOTUs (big black squares) forming viral clusters with Nonlabens phage P12024S and P12024L and with each other.

Protein-based clustering using vConTACT2 showed similarities of some South Georgia vOTUs to known marine phages. vOTU_07 shared similarities with Nonlabens phage P12024L (GenBank, <u>JQ823123</u>, Figure 4a, b), which originates from coastal seawater [72]. vOTU_36 and vOTU_37, detected in rainwater, formed VC_747_0 sharing protein similarities with the Baltic Sea derived Flavobacterium phage vB_FspP_elemoA_8-9C (GenBank, <u>MT497073</u>). Based on VipTree, vOTU_35 was related to the widespread oceanic Cyanophage S-TIM5 (GenBank, <u>NC_019516</u>) [73, 74] and estuarine Synechococcus phage S-CBWM1 (GenBank, <u>NC_048106</u>, Figure 5a) [75], vOTU_28, vOTU_29, vOTU_36, and vOTU_37 were related to different Flavobacterium phages of the genera *Elemovirus* and *Immutovirus* from the Baltic Sea [76], and vOTU_27 to different Cellulophaga phages isolated from the Baltic Sea [77] and North Sea [78]. In total, 15 vOTUs were related to viruses known to have marine hosts or marine origin themselves (Figure 4a, Table S4), providing evidence that the airborne viral community sampled here in South Georgia was influenced by marine input.

**Figure 5:**
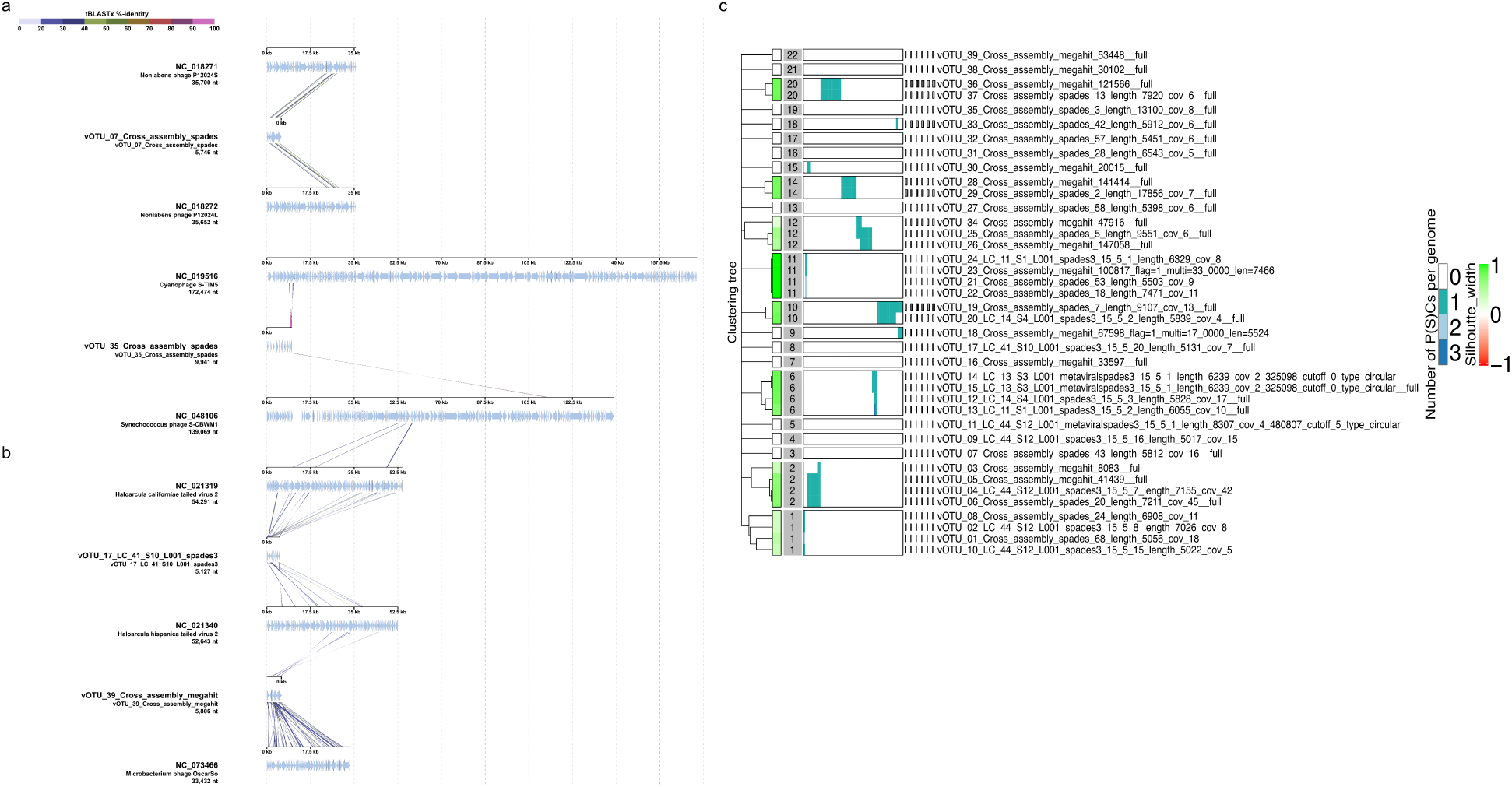
Sequence alignment and proteome clustering of annotated vOTUs. a) Sequence alignment of vOTUs with marine viruses. vOTU_07 demonstrate nucleotide identity with two Nonlabens phages. Sequence alignment of vOTU_35 indicate nucleotide identity with two marine cyanophages. b) Sequence alignment of vOTUs from extremophilic hosts. vOTU_17 demonstrate tBLASTx identity with sequences of two archaeal viruses. Sequence alignment of vOTU_39 shows tBLASTx identity with a high genomic GC content containing actinobacteriophage OscarSo (Genbank, <u>OP434449.1</u>) infecting *Microbacterium radiodurans*. c) Viral proteome clustering of all vOTUs with Virclust indicates eight different clades (clusters 1, 2, 6, 10-12, 14 and 20) out of the predicted 22.

Functional annotations of the 39 vOTUs are given in Table S5. vOTU_19, related to Nonlabens phages P12024S (GenBank, <u>NC_018271</u>) and P12024L (GenBank, <u>NC_018272</u>) (Figure 5a), had the most ORFs (19), encoding for proteins involved in host lytic machinery such as holin (ORF6) and peptidase (ORF9) [79]. Sequence alignment and tBLASTx analysis of the vOTUs indicated potential similarities with viruses from different ecosystems. Several protein coding genes of vOTU_07 showed ∼50% sequence identity with phages P12024S (GenBank, <u>NC_018271</u>) and P12024L (GenBank, <u>NC_018272</u>) (Figure 5a), which infect the marine bacterial host *Persicivirga* (family Flavobacteriaceae) [72]. Additionally, vOTU_17 and vOTU_39 displayed protein similarities with viruses that infect extremophilic hosts such as haloviruses HHTV-2 (GenBank, <u>NC_021340.1</u>) and HCTV-2 (GenBank, <u>NC_021319.1</u>) which infects the archaeon *Haloarcula californiae* isolated from salt brines (DSMZ, <u>DSM 8905</u>) [80] and phage OscarSo (GenBank, <u>NC_073466.1</u>), which infects *Microbacterium radiodurans* NRRL B-24799, a highly UV-tolerant bacterium [81], respectively (Figure 5b).

Hierarchical and phylogenetic clustering of viral core proteins highlighted clustering into eight distinct clades (clusters 1, 2, 6, 10-12, 14 and 20) out of the predicted 22 (Figure 5c, Table S6). These clusters in the hierarchical tree demonstrated protein overlaps and provided similar phylogenetic reconstructions as observed in the ViPtree analysis (Figure 5a, b). Clusters 2, 6, and 11 had the greatest number of members (four) while cluster 20, comprised of just two members, shared the greatest number of common core genes. This cluster is formed from vOTU_36 and vOTU_37 and was related to Flavobacterium phage vB_FspP_elemoA_8-9C (GenBank, <u>MT497073</u>) in vConTACT2. Overall, we found high diversity of the proteome in the analyzed vOTUs, with proteins overlapping among scaffolds being phylogenetically similar.

### Spatial distribution of selected protein homologs in the major oceans

To investigate the ecological significance and environmental distribution of viral proteins with potential functional roles in photosynthesis and UV protection, we analysed homologous sequences from vOTU_35 and vOTU_39 using a metagenomic database from the Tara Oceans project. VOTU_35 shared protein identity with the Cyanophage S-TIM5 (Figure 5a), which encoded the photosynthetic reaction centre protein D1 with a psbA domain (vOTU_35_ORF14) (Figure 6a). The D1 and D2 reaction centre proteins form a heterodimer responsible for the establishment of photosystem II in cyanobacteria [82, 83], which is the most light-sensitive complex of cyanobacteria [84]. Another protein from vOTU_39 shared similarity with a protein from the phage OscarSo (GenBank, <u>NC_073466.1</u>) that infects the extremophilic UV-tolerant host *Microbacterium radiodurans* [81]. The phage carries a gene encoding for the protein from the impB/mucB/samB family with a DinP domain having functions in the bacterial SOS (DNA damage) response and thus has relevance in UV protection (Pfam database, <u>PF00817</u>) [85] (Figure 6a). To understand the distribution of these protein sequences and the possible origin of the encoding vOTUs, we explored the biogeography of the protein sequences based on matches with the Ocean Gene Atlas. The global distribution of the two protein homologs shows distinct patterns in terms of geographic spread and relative abundance. The protein homolog of the photosynthetic reaction centre protein D1 is widely distributed across Tara Oceans stations, with notable concentrations across the South Atlantic Ocean, western Indian Ocean, and eastern tropical Atlantic, as well as in parts of the central South Pacific. The highest abundances are observed in central Africa and the Indian Ocean region. This homolog appears across multiple size fractions, including 0 – 0.22 μm, 0.1 – 0.22 μm, and 0.22 – 0.45 μm, and exhibits a relatively high abundance range from 9.94e-6 to 1.02e-3. In contrast, the protein homolog impB/mucB/samB with a DinP domain has a more restricted distribution, being primarily detected across the South Atlantic Ocean, the western Indian Ocean, and the eastern equatorial Pacific Ocean. It is present in the same particle size fractions as the other homolog but at much lower abundances, ranging from 6.68e-9 to 1.06e-5. The highest abundances of this homolog are localized to a single prominent site in the South Atlantic Ocean (station Tara_076).

**Figure 6:**
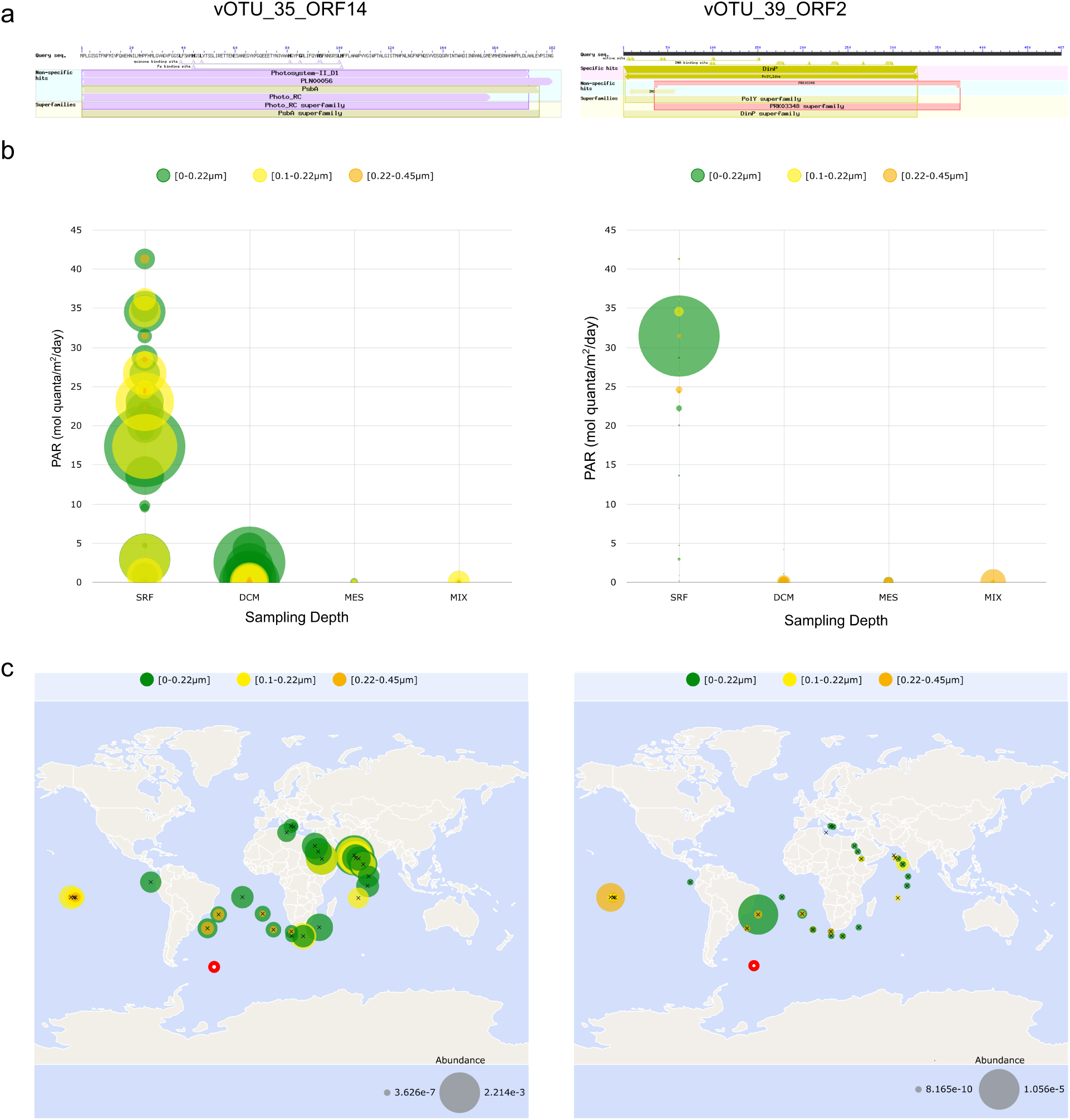
Distribution and abundance of viral protein sequence homologs in the major oceans. a) Domain analysis of photosynthetic reaction centre protein D1 derived from vOTU_35 and impB/mucB/samB derived from vOTU_39. b) the abundance of the protein homolog for the photosynthetic reaction centre protein D1 and impB/mucB/samB with a DinP domain in three different filtered fractions and across different sampling depths (SRF = surface water, DCM = deep chlorophyll maximum, MES = mesopelagic zone, MIX = mixed layer) and in relation to photosynthetically active radiation (PAR). c) Distribution of protein homologs across stations of the Tara Ocean expedition based on BLASTp to the Tara Oceans Microbiome Reference Genome Catalog v1 OM-RGC_v1 as obtained from Ocean Gene Atlas. Red circle indicates the position of South Georgia. Size of bubble indicates abundance in b and c.

Homologs of photosynthetic reaction centre protein D1 (vOTU_35_ORF14) were widely detected in several Tara Oceans sampling locations in the surface water layer (SRF) at varying PAR ranging from approximately 0.07 mol quanta m^−2^ day^−1^ to 41.3 mol quanta m^−2^ day^−1^ as compared to the mesopelagic zone (MES), where almost no PAR was observed, deep chlorophyll maximum (DCM) with PAR ranging from approximately 0.038 mol quanta m^−2^ day^−1^ to 2.52 mol quanta m^−2^ day^−1^, and the marine epipelagic mixed layer (MIX) with PAR ranging from approximately 0.00002 mol quanta m^−2^ day^−1^ to 0.000059 mol quanta m^−2^ day^−1^ (Figure 6b, Table S7). A total of 2625 hits were obtained for this protein, with an abundance value of 192236. Taxonomic distribution of these homologs indicated high prevalence of this protein in viral genomes (37%) (Figure S1). The UV protection related protein (vOTU_39_ORF2) was mostly abundant in the SRF as compared to the MES, DCM, and MIX layers, however, the distribution across the sampling locations was rather sparse as compared to vOTU_35_ORF14 (Figure 6b, Table S7&8). Only a total of 59 hits were obtained for this impB/mucB/samB family protein and the number of abundance measures was 1924, while for the photosynthetic reaction centre protein D1 we received 2625 hits and the number of abundance measures was 192236. Viral homologs of the impB/mucB/samB family protein could not be found. The closest homolog predicted by the Ocean Gene Atlas for this protein was the impB/mucB/samB and DinP domain-containing protein of the bacterium, *Vulcanimicrobium alpinum* (GenBank, <u>WP_317995911.1</u>) [86], isolated from a fumarole ice cave at high altitude on the volcano Mount Erebus in Victoria Land, Antarctica (Figure S2). Both protein homologs were present at sampling location TARA_076, being closer to the sampling site of our study than most other Tara Ocean stations (Figure 6c).

### Microbial community composition and host assignments

Only three hosts could be predicted for the 39 vOTUs using iPhoP. These were the genera *Rickettsia* (confidence score 90.8, phylum Pseudomonadota) for vOTU_34, *Myroides* for vOTU18 (confidence score 90.9, phylum Bacteroidota) and UBA9320 for vOTU_37 (confidence score 90.9, phylum Bacteroidota), the latter being a metagenome assembled genome from the Central Arctic [87]. Microbial community profiling (mainly prokaryotes, with a small number of reads of Basidiomycota) showed that the airborne community was dominated by Actinobacteria, Proteobacteria (now Pseudomonadota) and unassigned taxa. More unassigned taxa were found in the inland DMP samples (up to 80% relative abundance) than at the coastal KEP samples (up to 18% relative abundance, Table S9, Figure S3). BLASTing vOTUs to the IMG/VR database revealed hits to orders Ortervirales (vOTU_01, vOTU_23, vOTU_24) and Algavirales (vOTU_03), to the class Caudoviricetes (vOTU_07, vOTU_12 – vOTU_16, vOTU_18, vOTU_19, vOTU_20, vOTU_25 – vOTU_29, vOTU_33, vOTU_34 – vOTU_37, vOTU_39), and to the order Priklausovirales (vOTU_31). We acknowledge, however, that the distinction between viruses and retroelements can be challenging for Ortervirales-like sequences. Predicted hosts were of the phyla Actinobacteriota (vOTU_12 – vOTU_15, Alphaproteobacteria (vOTU_39), Saprospiraceae (vOTU_36 & vOTU_37) and Spirochaetota (vOTU_12 – vOTU_15). Most viral hits were to marine, aquatic, seawater ecosystems from various parts of the world (Table S10).

## Discussion

### Aerosolization of different viral groups, site effects, and biogeographic distribution

Our data show that the airborne viral community composition over South Georgia is influenced by the sampling site on a local scale (coastal vs inland) and the sampling device used. We found typical eukaryotic viral groups like NCLDV, which often are parasitized upon by smaller DNA viruses, known informally as virophage. Both virus types have been previously reported after sampling from 15 m depth in Marguerite Bay in the Southern Ocean close to the Antarctic continent [30] and to influence algal host-virus interactions in a meromictic Antarctic lake [88]. About half of all the vOTUs identified corresponded to phages. Our data confirm that viruses of various groups become airborne and form the first baseline data on airborne viral communities from South Georgia, along with evidence of day-to-day fluctuations, and of influence by the marine environment.

More vOTUs were assembled from and were present based on read mapping at the immediately coastal KEP site, suggesting aerosolization of local marine viruses. Several vOTUs shared proteins with known cyanophages and phage isolates infecting typical marine heterotrophic bacteria and, additionally, typical marine prokaryotes such as Thaumarchaeota (*Nitrosopumilus* sp.), were part of the microbial community (Figure S3). An earlier study in the Arctic explored viral distribution and adaptation at the air-sea interface and showed that virus dispersal across the Arctic might be facilitated by aerosolization of viruses residing in the sea-surface microlayer, the uppermost 1 mm surface layer in contact with the atmosphere [89]. Viruses are active in the Antarctic microlayer [90] from where they can be emitted into aerosols [91]. As our sampling sites were located on a remote Southern Ocean island far from other terrestrial influences, finding such small-scale local differences in marine contributions to the airborne viral community was not our initial expectation as, in absolute terms, both sites are physically close to the coast. High GC content vOTUs known to be present in air and rain samples from Swedish air [60] were detected on one sampling day in the dataset explored here. This was a day with particularly low wind speeds, sunny weather, and a wind direction of 238 degrees (west-south-west), the latter contrasting with all other sampling, where the wind direction was north-west to north-north-west (Table S1). These high GC viruses shared protein identities with Microbacterium phage OscarSo (37.5% identity in case of vOTU_39), which has a GC content of 69.2% (https://phagesdb.org/phages/OscarSo/), and with two archaeal viruses of the halophilic archaeon *Haloarcula* (e.g. 34.4% and 34.7% in case of vOTU_17). OscarSo has been isolated from *Microbacterium radiodurans*, which is highly resistant to UV radiation and originates from sand of the Gobi Desert [81]. Hence, our findings provide some evidence that airborne viruses with high GC content could have extremophilic hosts, allowing speculation that such viruses have a non-marine origin and stem from the higher atmosphere or terrestrial influences such as desert dust. The majority of marine prokaryote genomes have a GC content ranging between 30% and 50% [92, 93], and associated viruses typically mirror the GC content of their hosts [94, 95].

The distance between the two sampling sites DMP and KEP is approximately 2 km and, hence, our data overall confirm short-range dispersal of Antarctic viruses with bioaerosols, with likely influences from sea spray and wet precipitation as precipitation occurred during some of the sampling events (Table S1). Viral alpha diversity did not differ significantly between the two sites, whereas bacterial diversity showed a significant difference [32]. The marine origin of certain viruses and the spread of specific genes suggest that viruses are primarily sourced through atmospheric transfer over long distances. We speculate that viral communities are less influenced by a study site than bacteria, because of longer residence times and lower deposition velocities in air, since they are smaller [16, 17]. Therefore, they can be dispersed further and can become part of another remote community more easily.

Proteomic alignment and classification of the vOTUs was consistent with the results of the phylogenetic and clustering analyses. We found sequence identity at the amino acid level of some vOTUs with viruses of typical marine bacteria and archaea. Specifically, two different proteins encoded by vOTU_35_ORF14 and vOTU_39_ORF2 shared several homologs that are widely distributed in the surface water layer of the major oceans (Pacific, Atlantic, and Indian Ocean), supporting a probable marine origin. While speculative, the functionality of these proteins and their potential involvement in modulating a host’s metabolism may be important. Phages present in diverse ecological habitats must be highly adaptive; for instance, this could involve the placement of structural genes in regions of above-average GC content of the genome [96], encoding specialized proteins promoting host survival and extremotolerance, or reverting to lysogenic or pseudolysogenic life cycles [97, 98]. Typical traits of marine phages include augmentation of host metabolism or enhancing viral fitness by the acquisition and expression of auxiliary metabolic genes (AMGs) [99]. Considering that phage-derived psbA has been demonstrated to support host photosynthetic activity in surface water [100], and the potential of the impB/mucB/samB gene cluster to support microbial growth in harsh environments is known [101], we hypothesize that vOTU_35_ORF14 and vOTU_39_ORF2 could potentially be AMGs supporting the replication of these viruses in the marine and atmospheric environments, respectively. This could be appropriate and adaptive, as organisms in both environments face challenging conditions, for instance surface water in the open ocean is oligotrophic [102], and the atmosphere is prone to desiccation and solar and UV radiation effects [103].

### Methodological challenges of sampling air viruses for metagenomics

Several viruses remained unclustered by vConTACT2 or could only be clustered with other viruses obtained in this study, suggesting the presence of previously unknown viral diversity. In interpreting our results, it is important to acknowledge that many viruses remain unclassified and undocumented in public repositories. As a result, the vConTACT2 reference database used in this study may not fully capture the diversity of viral sequences present in our samples, potentially leading to the underrepresentation of novel or poorly characterized viral taxa. In addition, the overall viral biomass in these air samples was low, a common challenge in aeromicrobiology [104, 105]. As in a previous air virus study [106], this led us to lower the generally-applied 10 kb length cut-off for the analysis of vOTUs proposed by the standard viromics guidelines [47] to accept smaller, 5 kb, viral scaffolds to be used for viral presence/absence determination. Such a step might also be appropriate for studies in other low and ultra-low biomass systems such as clean rooms or space equipment [107, 108]. However, accepting smaller fragments can also increase uncertainties in various downstream predictions as well as the probability of false-positive predictions. Because airborne viruses are present in low abundance, the detection of a vOTU in a sample is more likely to be affected by limited coverage compared to high biomass ecosystems, making it difficult to accurately interpret overlapping communities and day-to-day variation in diversity.

The type of sampling device used here clearly influenced the viral findings. The dry Coriolis sampled few viruses (a single virus was assembled) and none were found by mapping reads from dry Coriolis samples to vOTUs retrieved from other samplers. According to the manufacturer’s information, the Coriolis Compact has a flow rate of 50 L min^−1^ and is described as a device capable of virus sampling, but this device is not mentioned in the academic literature on this subject. The wet Coriolis (Coriolis µ) has been used extensively for viral air sampling [109, 110]. It has a higher flow rate of up to 300 L min^−1^ and collects particles into a liquid and, thus, might be more efficient than the dry Coriolis. Additionally, the liquid might be more suitable for conserving the viral particles. Both Coriolis samplers have a size range of 500 nm – 10 µm for collected particles, which should exclude most viruses. As we have no information on the particle size of the sampled viruses here, we can only speculate whether small viruses were not sampled or were only collected if attached to larger particles. Further tests are required to thoroughly validate the devices used for viral sampling, including tests on the stability of viral nucleic acids on filters and in liquid, as degradation of nucleic acids is possible during long-term air sampling [111]. Instead of sampling on 0.2 µm pore-sized membranes or Sterivex filters, future studies of viruses collected into liquid could be iron-flocculated or subjected to ultrafiltration and processed as viromes, as done for aquatic samples [112].

A larger diversity of viruses was sampled with the rain collector. Rain and aerosol-derived viruses can be genomically very different [60], while airborne microbial communities also vary between precipitation types and with seasonal variations in Antarctica [11, 113]. Work by Jiang, et al. [106] supports that the atmosphere harbors a distinct viral community while also revealing habitat-specific viral assemblages and AMGs. Supporting previous findings [60], our work revealed VCs only found in the rainwater samples, which could indicate that they originate from a higher altitude. The collection of different, non-overlapping vOTUs by our two sampling devices suggests that using a funnel collector alongside an impinger air sampling system provides a more comprehensive representation of the airborne viral community, which can vary between aerosols and rain.

## Acknowledgements

We acknowledge use of the HPC cluster DRACO, with tools and database kindly provided by the VEO group of Bas Dutilh at the FSU Jena. We acknowledge technical assistance by Luke Cockerton for DNA extractions and by Bingli Chai for submitting the sequence data to SRA.

## Ethics approval and consent to participate

Not applicable.

## Consent for publication

All authors consent to the publication of the manuscript.

## Data Availability

Sequencing data are stored in NCBI’s SRA under the Bioproject PRJNA1107129, Biosample IDs: SAMN41160173 - SAMN41160182. The vOTUs are deposited at figshare under doi: 10.6084/m9.figshare.26536105 and in NCBI’s BankIt under accession numbers PV563644 - PV563682. Weather data are downloadable via https://legacy.bas.ac.uk/cgi-bin/metdb-form-2.pl?tabletouse=U_MET.GRYTVIKEN_AWS&complex=1&idmask=.....acct=u_met&pass=weather.

## Author Contribution Statement

RD: data curation, formal analysis, investigation, methodology, visualization, validation, writing original draft, writing review & editing; LM: field sampling, data acquisition, writing review & editing; DAP: conceptualization, field sampling, writing review & editing; PC: interpretation, writing review and editing; JR: conceptualization, formal analysis, funding acquisition, project administration, supervision, visualization, validation, writing original draft, writing review & editing

## List of Supporting Material Legends

**Figure S1: Protein homolog analysis of Photosynthetic reaction centre protein D1 (psbA) vOTU_35_ORF2.** Krona plot demonstrates the proportion of the homologs in viral genome isolates from the Tara Oceans Microbiome Reference Genome Catalog v1 OM-RGC_v1.

**Figure S2: Protein homolog analysis of impB/mucB/samB from vOTU_39.** Phylogeny of impB/mucB/samB from vOTU_39_02 showing close similarity and ancestry with the impB/mucB/samB with a DinP domain containing protein of the extremophile *Vulcanimicrobium alpinum* (edges highlighted in red). Black triangles indicate collapsed nodes.

**Figure S3:** Relative abundance of air-derived prokaryotes based on a) phylum and b) class. DC=dry Coriolis, Sea-lvl= sea level, WC=wet Coriolis. LC refers to an internal sample number.

Table S1: Metadata

Table S2: vOTUs_relabd

Table S3: vOTUs_infos

Table S4: vOTU_ecosystems

Table S5: vOTU_annotations

Table S6: Protein_VirClust

Table S7: OTU35ORF14_OGeneAtlas

Table S8: vOTU39_ORF2_OGeneAtlas

Table S9: mOTUs_relabd

Table S10: IMG/VR hits

## References

1. Malard LA, Sabacka M, Magiopoulos I, Mowlem M, Hodson A, Tranter M, et al. Spatial variability of Antarctic surface snow bacterial communities. Front Microbiol. 2019;10:461;

2. Pearce DA, Bridge PD, Hughes KA, Sattler B, Psenner R, Russell NJ. Microorganisms in the atmosphere over Antarctica. FEMS Microbiol Ecol. 2009;69(2):143–57;

3. Pretorius I, Schou WC, Richardson B, Ross SD, Withers TM, Schmale DG, 3rd, et al. In the wind: Invasive species travel along predictable atmospheric pathways. Ecol Appl. 2023;33(3):e2806;

4. Cary SC, McDonald IR, Barrett JE, Cowan DA. On the rocks: the microbiology of Antarctic Dry Valley soils. Nat Rev Microbiol. 2010;8(2):129–38;

5. Cavicchioli R. Microbial ecology of Antarctic aquatic systems. Nat Rev Microbiol. 2015;13(11):691–706;

6. Verleyen E, Van de Vijver B, Tytgat B, Pinseel E, Hodgson DA, Kopalová K, et al. Diatoms define a novel freshwater biogeography of the Antarctic. Ecography. 2021;44(4):548–60;

7. Vyverman W, Verleyen E, Wilmotte A, Hodgson DA, Willems A, Peeters K, et al. Evidence for widespread endemism among Antarctic micro-organisms. Polar Science. 2010;4(2):103–13;

8. Cao Y, Yu X, Ju F, Zhan H, Jiang B, Kang H, et al. Airborne bacterial community diversity, source and function along the Antarctic Coast. Sci Total Environ. 2021;765:142700;

9. Pearce DA, Hughes KA, Lachlan-Cope T, Harangozo SA, Jones AE. Biodiversity of air-borne microorganisms at Halley Station, Antarctica. Extremophiles. 2010;14(2):145–59;

10. King-Miaow K, Lee K, Maki T, LaCap-Bugler D, Archer SDJ. Airborne microorganisms in Antarctica: transport, survival and establishment. The Ecological Role of Micro-Organisms in the Antarctic Environment. 2019:163–96;

11. Malard LA, Avila-Jimenez ML, Schmale J, Cuthbertson L, Cockerton L, Pearce DA. Aerobiology over the Southern Ocean - Implications for bacterial colonization of Antarctica. Environ Int. 2022;169:107492;

12. Heinrichs ME, Piedade GJ, Popa O, Sommers P, Trubl G, Weissenbach J, et al. Breaking the ice: A review of phages in polar ecosystems. Methods Mol Biol. 2024;2738:31–71;

13. Hendrix RW, Smith MC, Burns RN, Ford ME, Hatfull GF. Evolutionary relationships among diverse bacteriophages and prophages: all the world’s a phage. Proc Natl Acad Sci U S A. 1999;96(5):2192–7;

14. Nga DDY, Nhung VH, Nhan NT, Hien TT. Study on the concentration, composition, and recovery rate of bacterial bioaerosols after rainfall in Ho Chi Minh City. Environ Monit Assess. 2024;196(3):295;

15. Vishwakarma YK, Mayank, Ram K, Gogoi MM, Banerjee T, Singh RS. Bioaerosol emissions from wastewater treatment process at urban environment and potential health impacts. J Environ Manage. 2024;361:121202;

16. Reche I, D’Orta G, Mladenov N, Winget DM, Suttle CA. Deposition rates of viruses and bacteria above the atmospheric boundary layer. ISME J. 2018;12(4):1154–62;

17. Alsante AN, Thornton DCO, Brooks SD. Ocean Aerobiology. Front Microbiol. 2021;12(3143):764178;

18. Quinn PK, Collins DB, Grassian VH, Prather KA, Bates TS. Chemistry and related properties of freshly emitted sea spray aerosol. Chem Rev. 2015;115(10):4383–99;

19. Trainic M, Koren I, Sharoni S, Frada M, Segev L, Rudich Y, et al. Infection dynamics of a bloom-forming alga and its virus determine airborne coccolith emission from seawater. iScience. 2018;6:327–35;

20. Kepner RL, Jr., Wharton RA, Jr., Suttle CA. Viruses in Antarctic lakes. Limnol Oceanogr. 1998;43(7):1754–61;

21. Luhtanen A-M, Eronen-Rasimus E, Oksanen HM, Tison J-L, Delille B, Dieckmann GS, et al. The first known virus isolates from Antarctic sea ice have complex infection patterns. FEMS Microbiol Ecol. 2018;94(4):fiy028;

22. Paterson H, Laybourn-Parry J. Antarctic sea ice viral dynamics over an annual cycle. Polar Biol. 2011;35(4):491–7;

23. Lopez-Simon J, Vila-Nistal M, Rosenova A, De Corte D, Baltar F, Martinez-Garcia M. Viruses under the Antarctic Ice Shelf are active and potentially involved in global nutrient cycles. Nat Commun. 2023;14(1):8295;

24. Zablocki O, van Zyl L, Adriaenssens EM, Rubagotti E, Tuffin M, Cary SC, et al. High-level diversity of tailed phages, eukaryote-associated viruses, and virophage-like elements in the metaviromes of Antarctic soils. Appl Environ Microbiol. 2014;80(22):6888–97;

25. Sommers P, Fontenele RS, Kringen T, Kraberger S, Porazinska DL, Darcy JL, et al. Single-stranded DNA viruses in Antarctic cryoconite holes. Viruses. 2019;11(11):1022;

26. Guixa-Boixereu N, Vaqué D, Gasol JM, Sánchez-Cámara J, Pedrós-Alió C. Viral distribution and activity in Antarctic waters. Deep Sea Res Part II Top Stud Oceanogr. 2002;49(4-5):827–45;

27. Liang Y, Bai X, Jiang Y, Wang M, He J, McMinn A. Distribution of marine viruses and their potential hosts in Prydz Bay and adjacent Southern Ocean, Antarctic. Polar Biol. 2015;39(2):365–78;

28. Rahlff J, Bornemann TLV, Lopatina A, Severinov K, Probst AJ. Host-associated phages disperse across the extraterrestrial analogue Antarctica. Appl Environ Microbiol. 2022;88(10):e0031522;

29. Lopatina A, Medvedeva S, Shmakov S, Logacheva MD, Krylenkov V, Severinov K. Metagenomic Analysis of Bacterial Communities of Antarctic Surface Snow. Front Microbiol. 2016;7:398;

30. Piedade GJ, Schon ME, Lood C, Fofanov MV, Wesdorp EM, Biggs TEG, et al. Seasonal dynamics and diversity of Antarctic marine viruses reveal a novel viral seascape. Nat Commun. 2024;15(1):9192;

31. Banyard AC, Bennison A, Byrne AMP, Reid SM, Lynton-Jenkins JG, Mollett B, et al. Detection and spread of high pathogenicity avian influenza virus H5N1 in the Antarctic Region. Nat Commun. 2024;15(1):7433;

32. Malard LA, Convey P, Pearce DA. Daily turnover of airborne bacterial communities in the sub-Antarctic. Environ Microbiome. 2025;20(1):91;

33. Bushnell B. BBTools software package. URL http://sourceforgenet/projects/bbmap. 2014:578:579;

34. Joshi N, Fass J. Sickle: A sliding-window, adaptive, quality-based trimming tool for FastQ files. 2011;

35. Milanese A, Mende DR, Paoli L, Salazar G, Ruscheweyh HJ, Cuenca M, et al. Microbial abundance, activity and population genomic profiling with mOTUs2. Nat Commun. 2019;10(1):1014;

36. Ruscheweyh HJ, Milanese A, Paoli L, Karcher N, Clayssen Q, Keller MI, et al. Cultivation-independent genomes greatly expand taxonomic-profiling capabilities of mOTUs across various environments. Microbiome. 2022;10(1):212;

37. Nurk S, Meleshko D, Korobeynikov A, Pevzner PA. metaSPAdes: a new versatile metagenomic assembler. Genome Res. 2017;27(5):824–34;

38. Antipov D, Raiko M, Lapidus A, Pevzner PA. Metaviral SPAdes: assembly of viruses from metagenomic data. Bioinformatics. 2020;36(14):4126–9;

39. Kieft K, Zhou Z, Anantharaman K. VIBRANT: automated recovery, annotation and curation of microbial viruses, and evaluation of viral community function from genomic sequences. Microbiome. 2020;8(1):90;

40. Guo J, Bolduc B, Zayed AA, Varsani A, Dominguez-Huerta G, Delmont TO, et al. VirSorter2: a multi-classifier, expert-guided approach to detect diverse DNA and RNA viruses. Microbiome. 2021;9(1):37;

41. Camargo AP, Roux S, Schulz F, Babinski M, Xu Y, Hu B, et al. Identification of mobile genetic elements with geNomad. Nat Biotechnol. 2023;

42. Li D, Liu CM, Luo R, Sadakane K, Lam TW. MEGAHIT: an ultra-fast single-node solution for large and complex metagenomics assembly via succinct de Bruijn graph. Bioinformatics. 2015;31(10):1674–6;

43. Moraru C, Varsani A, Kropinski AM. VIRIDIC-A novel tool to calculate the intergenomic similarities of prokaryote-infecting viruses. Viruses. 2020;12(11);

44. Nayfach S, Camargo AP, Schulz F, Eloe-Fadrosh E, Roux S, Kyrpides NC. CheckV assesses the quality and completeness of metagenome-assembled viral genomes. Nat Biotechnol. 2021;39(5):578–85;

45. Langmead B, Salzberg SL. Fast gapped-read alignment with Bowtie 2. Nat Methods. 2012;9(4):357–9;

46. Nilsson E, Bayfield OW, Lundin D, Antson AA, Holmfeldt K. Diversity and host interactions among virulent and temperate Baltic Sea Flavobacterium phages. Viruses. 2020;12(2):158;

47. Roux S, Emerson JB, Eloe-Fadrosh EA, Sullivan MB. Benchmarking viromics: an in silico evaluation of metagenome-enabled estimates of viral community composition and diversity. PeerJ. 2017;5:e3817;

48. Bornemann TLV, Esser SP, Stach TL, Burg T, Probst AJ. uBin: A manual refining tool for genomes from metagenomes. Environ Microbiol. 2023;25(6):1077–83;

49. Hyatt D, Chen GL, Locascio PF, Land ML, Larimer FW, Hauser LJ. Prodigal: prokaryotic gene recognition and translation initiation site identification. BMC Bioinformatics. 2010;11(1):119;

50. Bolduc B, Jang HB, Doulcier G, You ZQ, Roux S, Sullivan MB. vConTACT: an iVirus tool to classify double-stranded DNA viruses that infect Archaea and Bacteria. PeerJ. 2017;5:e3243;

51. Cook R, Brown N, Redgwell T, Rihtman B, Barnes M, Clokie M, et al. INfrastructure for a PHAge REference Database: Identification of large-scale biases in the current collection of cultured phage genomes. Phage (New Rochelle). 2021;2(4):214–23;

52. Pandolfo M, Telatin A, Lazzari G, Adriaenssens EM, Vitulo N. MetaPhage: an automated pipeline for analyzing, annotating, and classifying bacteriophages in metagenomics sequencing data. mSystems. 2022;7(5):e0074122;

53. Shannon P, Markiel A, Ozier O, Baliga NS, Wang JT, Ramage D, et al. Cytoscape: a software environment for integrated models of biomolecular interaction networks. Genome Res. 2003;13(11):2498–504;

54. Shang J, Tang X, Guo R, Sun Y. Accurate identification of bacteriophages from metagenomic data using Transformer. Brief Bioinform. 2022;23(4):bbac258;

55. Shang J, Jiang J, Sun Y. Bacteriophage classification for assembled contigs using graph convolutional network. Bioinformatics. 2021;37(Suppl_1):i25–i33;

56. Shang J, Peng C, Liao H, Tang X, Sun Y. PhaBOX: a web server for identifying and characterizing phage contigs in metagenomic data. Bioinform Adv. 2023;3(1):vbad101;

57. Moraru C. VirClust-A tool for hierarchical clustering, core protein detection and annotation of (prokaryotic) viruses. Viruses. 2023;15(4):1007;

58. Nishimura Y, Yoshida T, Kuronishi M, Uehara H, Ogata H, Goto S. ViPTree: the viral proteomic tree server. Bioinformatics. 2017;33(15):2379–80;

59. Roux S, Fischer MG, Hackl T, Katz LA, Schulz F, Yutin N. Updated virophage taxonomy and distinction from polinton-like viruses. Biomolecules. 2023;13(2):204;

60. Rahlff J, Esser SP, Plewka J, Heinrichs ME, Soares A, Scarchilli C, et al. Marine viruses disperse bidirectionally along the natural water cycle. Nat Commun. 2023;14(1):6354;

61. Rice P, Longden I, Bleasby A. EMBOSS: the European Molecular Biology Open Software Suite. Trends Genet. 2000;16(6):276–7;

62. Shaffer M, Borton MA, McGivern BB, Zayed AA, La Rosa SL, Solden LM, et al. DRAM for distilling microbial metabolism to automate the curation of microbiome function. Nucleic Acids Res. 2020;48(16):8883–900;

63. Camargo AP, Nayfach S, Chen IA, Palaniappan K, Ratner A, Chu K, et al. IMG/VR v4: an expanded database of uncultivated virus genomes within a framework of extensive functional, taxonomic, and ecological metadata. Nucleic Acids Res. 2023;51(D1):D733–D43;

64. Marchler-Bauer A, Derbyshire MK, Gonzales NR, Lu S, Chitsaz F, Geer LY, et al. CDD: NCBI’s conserved domain database. Nucleic Acids Res. 2015;43(Database issue):D222–6;

65. Vernette C, Lecubin J, Sanchez P, Tara Oceans C, Sunagawa S, Delmont TO, et al. The Ocean Gene Atlas v2.0: online exploration of the biogeography and phylogeny of plankton genes. Nucleic Acids Res. 2022;50(W1):W516–W26;

66. Villar E, Vannier T, Vernette C, Lescot M, Cuenca M, Alexandre A, et al. The Ocean Gene Atlas: exploring the biogeography of plankton genes online. Nucleic Acids Res. 2018;46(W1):W289–W95;

67. Escobedo JF, Gomes EN, Oliveira AP, Soares J. Ratios of UV, PAR and NIR components to global solar radiation measured at Botucatu site in Brazil. Renewable Energy. 2011;36(1):169–78;

68. Letunic I, Bork P. Interactive Tree of Life (iTOL) v6: recent updates to the phylogenetic tree display and annotation tool. Nucleic Acids Res. 2024;52(W1):W78–W82;

69. Roux S, Camargo AP, Coutinho FH, Dabdoub SM, Dutilh BE, Nayfach S, et al. iPHoP: An integrated machine learning framework to maximize host prediction for metagenome-derived viruses of archaea and bacteria. PLoS Biol. 2023;21(4):e3002083;

70. McMurdie PJ, Holmes S. phyloseq: an R package for reproducible interactive analysis and graphics of microbiome census data. PLoS One. 2013;8(4):e61217;

71. Team RC. R: A language and environment for statistical computing, R Foundation for Statistical Computing, Vienna, Austria. URL https://www.R-project.org/. 2019;

72. Kang I, Jang H, Cho JC. Complete genome sequences of two *Persicivirga* bacteriophages, P12024S and P12024L. J Virol. 2012;86(16):8907–8;

73. Sabehi G, Shaulov L, Silver DH, Yanai I, Harel A, Lindell D. A novel lineage of myoviruses infecting cyanobacteria is widespread in the oceans. Proc Natl Acad Sci U S A. 2012;109(6):2037–42;

74. Baran N, Carlson MCG, Sabehi G, Peleg M, Kondratyeva K, Pekarski I, et al. Widespread yet persistent low abundance of TIM5-like cyanophages in the oceans. Environ Microbiol. 2022;24(12):6476–92;

75. Xu Y, Zhang R, Wang N, Cai L, Tong Y, Sun Q, et al. Novel phage–host interactions and evolution as revealed by a cyanomyovirus isolated from an estuarine environment. Environ Microbiol. 2018;20(8):2974–89;

76. Hoetzinger M, Nilsson E, Arabi R, Osbeck CMG, Pontiller B, Hutinet G, et al. Dynamics of Baltic Sea phages driven by environmental changes. Environ Microbiol. 2021;23(8):4576–94;

77. Holmfeldt K, Solonenko N, Shah M, Corrier K, Riemann L, Verberkmoes NC, et al. Twelve previously unknown phage genera are ubiquitous in global oceans. Proc Natl Acad Sci U S A. 2013;110(31):12798–803;

78. Bartlau N, Wichels A, Krohne G, Adriaenssens EM, Heins A, Fuchs BM, et al. Highly diverse flavobacterial phages isolated from North Sea spring blooms. ISME J. 2022;16(2):555–68;

79. Shah S, Das R, Chavan B, Bajpai U, Hanif S, Ahmed S. Beyond antibiotics: phage-encoded lysins against Gram-negative pathogens. Front Microbiol. 2023;14:1170418;

80. Sencilo A, Jacobs-Sera D, Russell DA, Ko CC, Bowman CA, Atanasova NS, et al. Snapshot of haloarchaeal tailed virus genomes. RNA Biol. 2013;10(5):803–16;

81. Zhang W, Zhu HH, Yuan M, Yao Q, Tang R, Lin M, et al. *Microbacterium radiodurans* sp. nov., a UV radiation-resistant bacterium isolated from soil. Int J Syst Evol Microbiol. 2010;60(Pt 11):2665–70;

82. Shi LX, Schroder WP. The low molecular mass subunits of the photosynthetic supracomplex, photosystem II. Biochim Biophys Acta. 2004;1608(2-3):75–96;

83. Kamiya N, Shen JR. Crystal structure of oxygen-evolving photosystem II from *Thermosynechococcus vulcanus* at 3.7-A resolution. Proc Natl Acad Sci U S A. 2003;100(1):98–103;

84. Pathak J, Ahmed H, Singh PR, Singh SP, Häder D-P, Sinha RP. Chapter 7 - Mechanisms of Photoprotection in Cyanobacteria. In: Mishra AK, Tiwari DN, Rai AN, editors. Cyanobacteria. Academic Press; 2019. p. 145–71.

85. Smith BT, Walker GC. Mutagenesis and more: umuDC and the Escherichia coli SOS response. Genetics. 1998;148(4):1599–610;

86. Yabe S, Muto K, Abe K, Yokota A, Staudigel H, Tebo BM. *Vulcanimicrobium alpinus* gen. nov. sp. nov., the first cultivated representative of the candidate phylum “Eremiobacterota”, is a metabolically versatile aerobic anoxygenic phototroph. ISME Commun. 2022;2(1):120;

87. Winder JC, Boulton W, Salamov A, Eggers SL, Metfies K, Moulton V, et al. Genetic and structural diversity of prokaryotic ice-binding proteins from the Central Arctic Ocean. Genes (Basel). 2023;14(2);

88. Yau S, Lauro FM, DeMaere MZ, Brown MV, Thomas T, Raftery MJ, et al. Virophage control of antarctic algal host-virus dynamics. Proc Natl Acad Sci U S A. 2011;108(15):6163–8;

89. Rahlff J, Westmeijer G, Weissenbach J, Antson A, Holmfeldt K. Surface microlayer-mediated virome dissemination in the Central Arctic. Microbiome. 2024;12(1):218;

90. Vaqué D, Boras JA, Arrieta JM, Agusti S, Duarte CM, Sala MM. Enhanced viral activity in the surface microlayer of the Arctic and Antarctic Oceans. Microorganisms. 2021;9(2):317;

91. Aller JY, Kuznetsova MR, Jahns CJ, Kemp PF. The sea surface microlayer as a source of viral and bacterial enrichment in marine aerosols. J Aerosol Sci 2005;36(5-6):801–12;

92. Luo H, Thompson LR, Stingl U, Hughes AL. Selection maintains low genomic GC content in marine SAR11 lineages. Mol Biol Evol. 2015;32(10):2738–48;

93. Yan W, Feng X, Zhang W, Nawaz MZ, Luo T, Zhang R, et al. Genomes of diverse isolates of *Prochlorococcus* high-light-adapted clade II in the Western Pacific Ocean. Front Mar Sci. 2021;7:619826;

94. Bahir I, Fromer M, Prat Y, Linial M. Viral adaptation to host: a proteome-based analysis of codon usage and amino acid preferences. Mol Syst Biol. 2009;5(1):311;

95. Simon D, Cristina J, Musto H. Nucleotide composition and codon usage across viruses and their respective hosts. Front Microbiol. 2021;12:646300;

96. Das R, Rahlff J. Phage genome architecture and GC content: Structural genes and where to find them. bioRxiv. 2024:2024.06.05.597531;

97. Irby I, Broddrick JT. Microbial adaptation to spaceflight is correlated with bacteriophage-encoded functions. Nat Commun. 2024;15(1):3474;

98. Hwang Y, Rahlff J, Schulze-Makuch D, Schloter M, Probst AJ. Diverse viruses carrying genes for microbial extremotolerance in the Atacama Desert hyperarid soil. mSystems. 2021;6(3):e00385–21;

99. Heyerhoff B, Engelen B, Bunse C. Auxiliary metabolic gene functions in pelagic and benthic viruses of the Baltic Sea. Front Microbiol. 2022;13:863620;

100. Sharon I, Tzahor S, Williamson S, Shmoish M, Man-Aharonovich D, Rusch DB, et al. Viral photosynthetic reaction center genes and transcripts in the marine environment. ISME J. 2007;1(6):492–501;

101. Wang SY, Chen YP, Huang RF, Wu YL, Ho ST, Li KY, et al. Subspecies classification and comparative genomic analysis of *Lactobacillus kefiranofaciens* HL1 and M1 for potential niche-specific genes and pathways. Microorganisms. 2022;10(8):1637;

102. Goldman JC. Oceanic nutrient cycles. In: Fasham MYR, editor. Flows of Energy and Materials in Marine Ecosystems: Theory and Practice. Springer New York, NY: Springer; 1984. p. 137–70.

103. Polymenakou PN. Atmosphere: A source of pathogenic or beneficial microbes? Atmosphere. 2012;3(1):87–102;

104. Hou J, Fujiyoshi S, Perera IU, Nishiuchi Y, Nakajima M, Ogura D, et al. Perspectives on sampling and new generation sequencing methods for low-biomass bioaerosols in atmospheric environments. J Indian Inst Sci. 2023;103(3):1–11;

105. Harnpicharnchai P, Pumkaeo P, Siriarchawatana P, Likhitrattanapisal S, Mayteeworakoon S, Ingsrisawang L, et al. AirDNA sampler: An efficient and simple device enabling high-yield, high-quality airborne environment DNA for metagenomic applications. PLoS One. 2023;18(6):e0287567;

106. Jiang T, Guo C, Yu H, Wang Z, Zheng K, Zhang X, et al. Habitat-Dependent DNA viral communities in atmospheric aerosols: Insights from terrestrial and marine ecosystems in East Asia. Environ Int. 2025;197:109359;

107. Highlander SK, Wood JM, Gillece JD, Folkerts M, Fofanov V, Furstenau T, et al. Multi-faceted metagenomic analysis of spacecraft associated surfaces reveal planetary protection relevant microbial composition. PLoS One. 2023;18(3):e0282428;

108. Zhang Y, Xin CX, Zhang LT, Deng YL, Wang X, Chen XY, et al. Detection of fungi from low-biomass spacecraft assembly clean room aerosols. Astrobiology. 2018;18(12):1585–93;

109. Verreault D, Gendron L, Rousseau GM, Veillette M, Masse D, Lindsley WG, et al. Detection of airborne lactococcal bacteriophages in cheese manufacturing plants. Appl Environ Microbiol. 2011;77(2):491–7;

110. Letourneau V, Gagne MJ, Vyskocil JM, Brochu V, Robitaille K, Gauthier M, et al. Hunting for a viral proxy in bioaerosols of swine buildings using molecular detection and metagenomics. J Environ Sci (China). 2025;148:69–78;

111. Boifot KO, Skogan G, Dybwad M. Sampling efficiency and nucleic acid stability during long-term sampling with different bioaerosol samplers. Environ Monit Assess. 2024;196(6):577;

112. Langenfeld K, Chin K, Roy A, Wigginton K, Duhaime MB. Comparison of ultrafiltration and iron chloride flocculation in the preparation of aquatic viromes from contrasting sample types. PeerJ. 2021;9:e11111;

113. Els N, Larose C, Baumann-Stanzer K, Tignat-Perrier R, Keuschnig C, Vogel TM, et al. Microbial composition in seasonal time series of free tropospheric air and precipitation reveals community separation. Aerobiologia. 2019;35:671–701;

114. Weatherall P, Ferreras SC, Cardigos SD, Cornish N, Davidson SR, Dorschel B, et al. The GEBCO_2024 Grid-a continuous terrain model of the global oceans and land. 2024.

