## Supplement Material for "Spatio-temporal variability of airborne viruses over sub-Antarctic South Georgia: Influence of sampling methodology and proximity of the ocean"

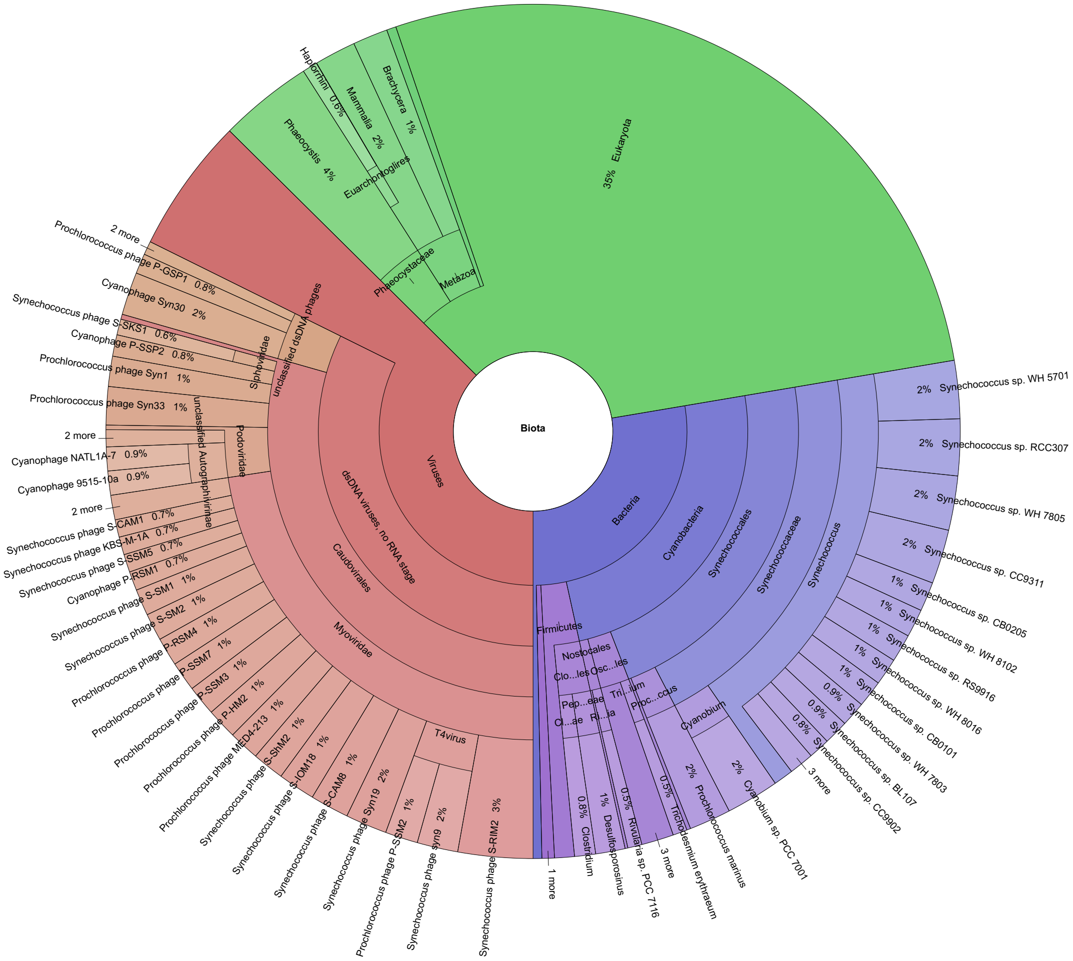


**Figure S1: Protein homolog analysis of Photosynthetic reaction centre protein D1 (psbA) vOTU_35_ORF2.** Krona plot demonstrates the proportion of the homologs in viral genome isolates from the Tara Oceans Microbiome Reference Genome Catalog v1 OM-RGC_v1.

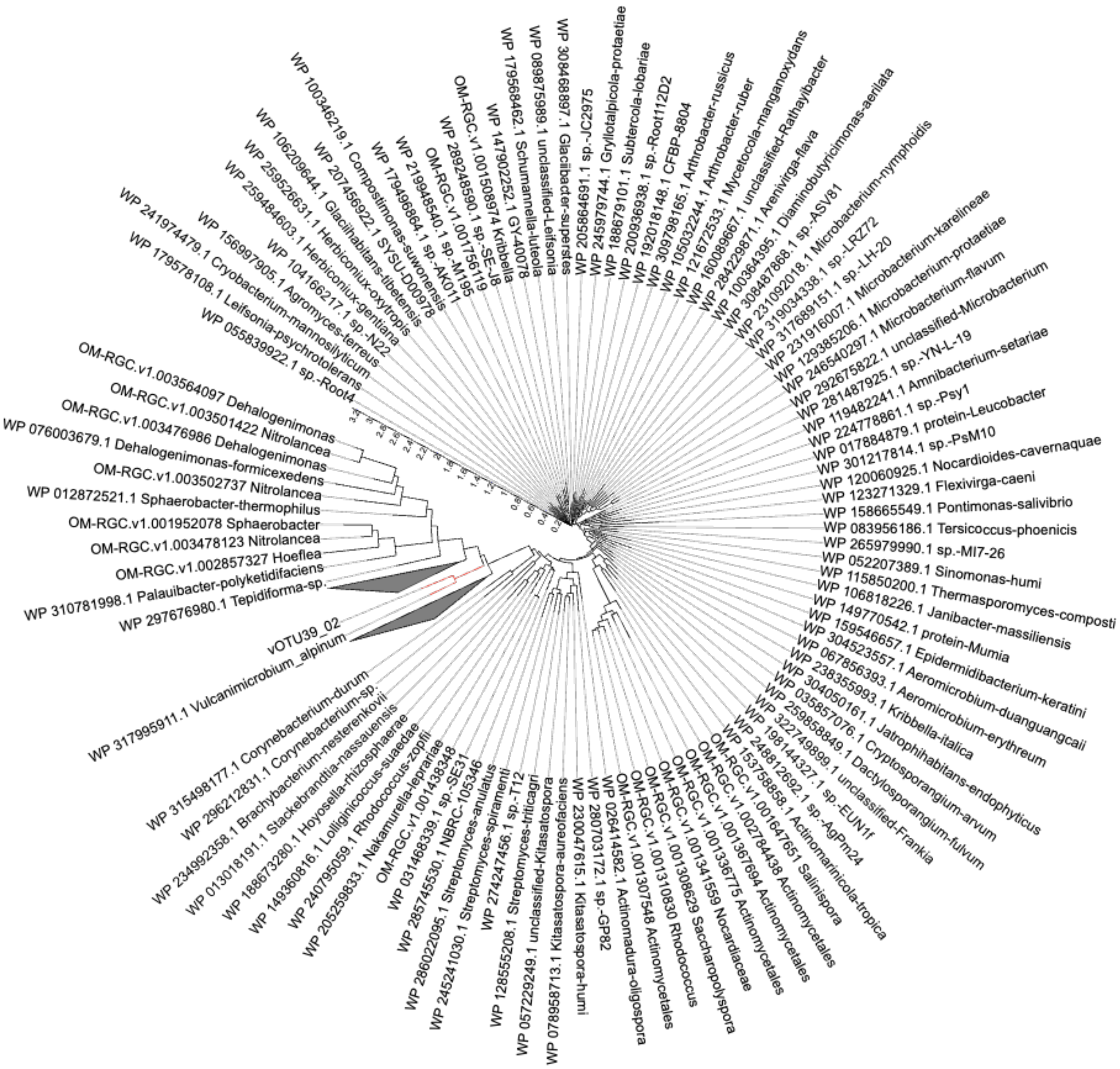


**Figure S2: Protein homolog analysis of impB/mucB/samB from vOTU_39.** Phylogeny of impB/mucB/samB from vOTU_39_02 showing close similarity and ancestry with the impB/mucB/samB with a DinP domain containing protein of the extremophile *Vulcanimicrobium alpinum* (edges highlighted in red). Black triangles indicate collapsed nodes.

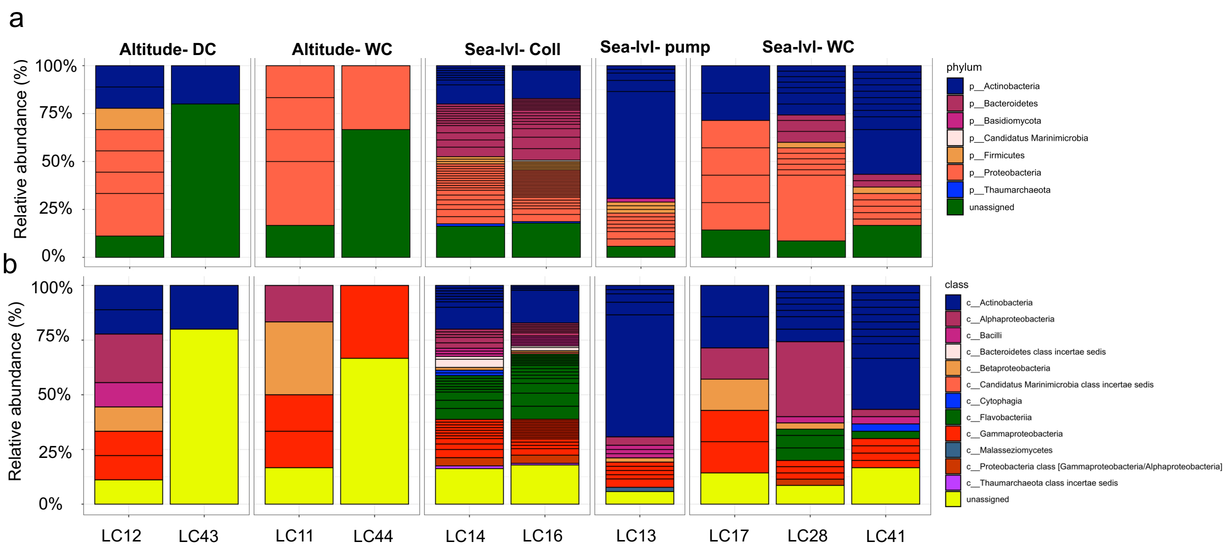


**Figure S3:** Relative abundance of air-derived prokaryotes based on a) phylum and b) class. DC=dry Coriolis, Sea-lvl= sea level, WC=wet Coriolis. LC refers to an internal sample number.
